# Comparison of village dog and wolf genomes highlights the pivotal role of the neural crest in dog domestication

**DOI:** 10.1101/118794

**Authors:** Amanda L. Pendleton, Feichen Shen, Angela M. Taravella, Sarah Emery, Krishna R. Veeramah, Adam R. Boyko, Jeffrey M. Kidd

**Affiliations:** Department of Human Genetics, University of Michigan, Ann Arbor, MI, 48109 USA.; Department of Ecology and Evolution, Stony Brook University, Stony Brook, NY 11794,USA.; Department of Biomedical Sciences, Cornell University, Ithaca, New York, 14853 USA.; Department of Computational Medicine and Bioinformatics, University of Michigan, Ann Arbor, MI 48109 USA.

**Keywords:** domestication, canine, selection scan, neural crest, retinoic acid

## Abstract

**Background:** Dogs (*Canis lupus familiaris*) were domesticated from gray wolves between 10-40 kya in Eurasia, yet details surrounding the process of domestication remain unclear. The vast array of phenotypes exhibited by dogs mirror other domesticated animal species, a phenomenon known as the Domestication Syndrome. Here, we use signatures persisting in the dog genome to identify genes and pathways altered by the intensive selective pressures of domestication.

**Results:** We identified 246 candidate domestication regions containing 10.8Mb of genome sequence and 178 genes through whole-genome SNP analysis of 43 globally distributed village dogs and 10 wolves. Comparisons with ancient dog genomes suggest that these regions reflect signatures of domestication rather than breed formation. The strongest hit is located in the *Retinoic Acid-Induced 1* (*RAI1*) gene, mutations of which cause Smith-Magenis syndrome. The identified regions contain a significant enrichment of genes linked to neural crest cell migration, differentiation and development. Read depth analysis suggests that copy number variation played a minor role in dog domestication.

**Conclusion:** Our results indicate that phenotypes distinguishing domesticated dogs from wolves, such as tameness, smaller jaws, floppy ears, and diminished craniofacial development, are determined by genes which act early in embryogenesis. These differences are all phenotypes of the Domestication Syndrome, which can be explained by decreases in neural crest cells at these sites. We propose that initial selection during early dog domestication was for behavior, a trait also influenced by genes which act in the neural crest, which secondarily gave rise to the phenotypes of modern dogs.

## Background

The process of animal domestication by humans was complex and multi-staged, resulting in the disparate appearances and behaviors of domesticates relative to their wild ancestors [1–3]. In 1868, Darwin noted that numerous traits are shared among domesticated animal species, an observation that has since been classified as the “Domestication Syndrome” (DS) [4]. DS is a phenomenon where diverse phenotypes are shared among phylogenetically distinct domesticated species but absent in their wild ancestors. Such traits include increased tameness, shorter muzzles/snouts, smaller teeth, more frequent estrous cycles, floppy ears, reduced brain size, depigmentation of skin or fur, and loss of hair.

During the domestication process, the most desired traits are subject to selection. This selection process may result in detectable genetic signatures such as alterations in allele frequencies [5–11], amino acid substitution patterns [12–14], and linkage disequilibrium patterns [15,16]. Though numerous genome selection scans have been performed within a variety of domesticated animal taxa [5–11,17], no single “domestication gene” shared across domesticates has been identified [18,19]. However, in the context of DS, this is not unexpected given more than a dozen diverse behavioral and complex physical traits fall under the syndrome. Based on the specific range of traits associated with DS, it is likely that numerous genes with pleiotropic effects likely contribute through mechanisms which act early in organismal development [18,19]. For this reason, the putative role of the neural crest in domestication has gained traction. While embryonic neural crest cells are the progenitors of most of the adult vertebrate body, these cells also influence behavior. Functions of the adrenal and pituitary systems, also derived from neural crest cells, influence aggression and the “fight or flight” behavioral reactions, and have found to be differentially affected in domesticates [20].

No domestic animal has shared more of its evolutionary history in direct contact with humans than the dog (*Canis lupus familiaris*), living alongside humans for tens of thousands of years since domestication from its ancestor the gray wolf (*Canis lupus*).

Despite numerous studies, vigorous debate still persist regarding the location, timing, and number of dog domestication events [21–25]. Several studies using related approaches have attempted to identify genomic regions which are highly differentiated between dogs and wolves, with the goal of identifying candidate targets of selection during domestications (candidate domestication regions, CDRs). In these studies breed dogs either fully or partially represented dog genetic diversity [24,26–29]. Most modern breeds arose ~300 years ago [30] and thus contain only a small portion of the genetic diversity found among the vast majority of extant dogs. Instead, semi-feral village dogs are the most abundant and genetically diverse modern dog populations, and have undergone limited targeted selection by humans since initial domestication [22,31]. These two dog groups represent products of two severe bottlenecks in the evolution of the domestic dog, the first resulting from the initial domestication of gray wolves, and the second from modern breed formation [32,33]. Studies based on breed dogs may therefore confound signatures associated with these two events. Indeed, we recently reported [34] that neither ancient nor modern village dogs could be genetically distinguished from wolves at 18 of 30 previously identified CDRs [26,27]. Furthermore, most of these studies employed empirical outlier approaches wherein the extreme tail of differentiated loci are assumed to differ due to the action of selection [35]. Freedman et al. [29], extended these studies through the use of a simulated demographic history to identify loci whose variability is unlikely to result from a neutral population history of bottlenecks and migration.

In this study, we reassess candidate domestication regions in dogs using genome sequence data from a globally diverse collection of village dogs and wolves. First, using methods previously applied to breed dog samples, we show that the use of semi-feral village dogs better captures dog genetic diversity and identifies loci more likely to be truly associated with domestication. Next, we perform a scan for CDRs in village dogs utilizing the XP-CLR statistic, refine our results by including haplotypes from ancient dogs, and present a revised set of pathways altered during dog domestication. Finally, we perform a scan for copy-number differences between village dogs and wolves, and identify additional copy-number variation at the starch-metabolizing gene *amylase-2b* (*AMY2b*) that is independent of the *AMY2b* tandem expansion previously found in dogs [26,36–38].

## Results

### Use of village dogs eliminates bias in domestication scans associated with breed formation

#### Comparison using F ST outlier approaches

We adapted methods from previous selection scan studies in dogs to determine the impact of sample choice (i.e. breed versus village dogs) on the detection of selective sweeps associated with early domestication pressures. We employed sliding window selection scan methodologies similar to those implemented in [26,27] to localize genomic intervals with unusual levels of allele frequency differentiation between the village dog and wolf populations. First, through ADMIXTURE [39] and identity-by-state (IBS) analyses, we identified a collection of 43 village dog and 10 gray wolf unrelated samples (Figure 1a) that have less than 5% dog-wolf admixed ancestry (see Methods). Principal component analysis illustrates the genetic separation between village dogs and wolves, and largely reflects the geographic distribution of these populations (Figure 1b and 1c). Next, we calculated average F_ST_ values in overlapping 200 kb sliding windows with a step-size of 50 kb across the genome. As in [26,27], we performed a Z-transformation to normalize the resulting values and identified windows with a ZF_ST_ score greater than 5 (autosomes) or 3 (X chromosome) as candidate domestication regions. Following merging, this outlier procedure identified 31 CDRs encompassing 12.3 Mb of sequence (Additional File 1: Table S1). As in previously studies, a 550 kb region on chromosome 6 (46.80-47.35 Mb) that contains the *pancreatic amylase 2B* (*AMY2B*) and *RNA Binding Region Containing 3* (*RNPC3*) genes had the highest observed average ZF_ST_ score (ZF_ST_ = 7.67).

**Figure 1.**
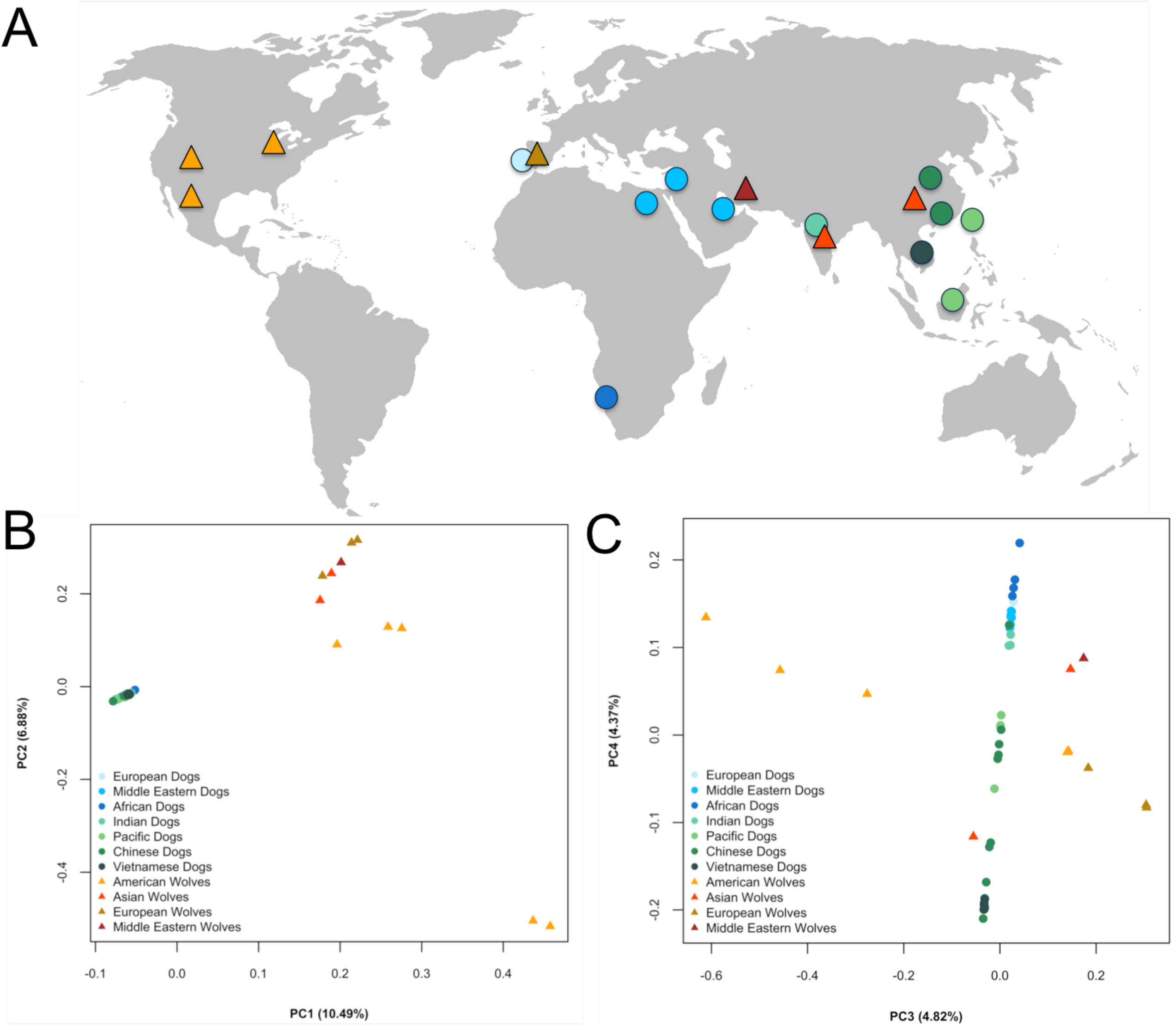
Origin and diversity of sampled village dogs and wolves. (A) The approximate geographic origin of the village dog (circles) and gray wolf (triangles) genome samples included in our analysis. Results of a Principal Component Analysis of the filtered village dog (N=43) and gray wolf set (N=11) are shown. Results are projected on (B) PC1 and PC2 and (C) PC3 and PC4. Colors in all figures correspond to sample origins and are explained in the PCA legends.

Only 15 of these 31 regions intersect with those reported in [26] and [27] (Figure 2a). To further explore this discrepancy, we visually assessed whether the dog or wolf haplotype is present at the loci reported in these earlier studies in 46 additional canine samples, including three ancient European dogs ranging in age from 5,000-7000 years old (see Methods; [21,34]). Likely due to the absence of village dogs in their study, some Axelsson loci appear to contain selective sweeps associated with breed formation, as evidenced by the presence of the wild haplotype in ancient and village dogs (example in Figure 2b). Although all autosomal sweeps identified by [8] intersected with CDRs from our study, seven X chromosome Cagan and Blass windows did not meet the thresholds of significance from our SNP sets (example in Additional File 2: Figure S1). We performed F_ST_ scans and Z transformations for windows on autosomes and X chromosome together, which may falsely inflate F_ST_ signals on the X due to smaller effective population sizes and correspondingly higher expected levels of genetic drift on the X chromosome. More detailed analysis of the loci highlighted in these two studies will be elaborated in the following section.

**Figure 2.**
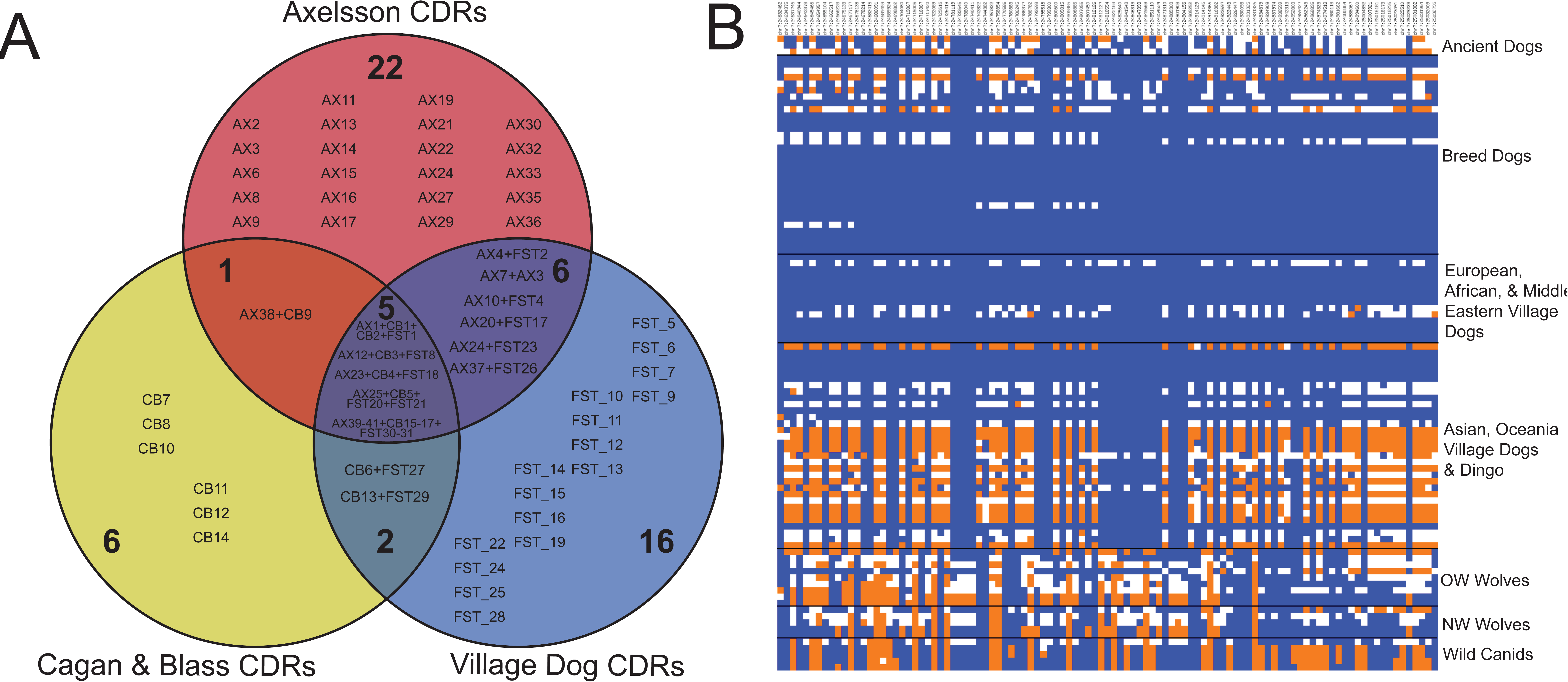
Comparison with previously published candidate domestication regions. (A) Venn diagram of intersecting village dog, Axelsson et al. 2013 (AX), and Cagan and Blass 2016 (CB) CDRs. (B) Genotype matrix for 130 SNPs within chr7: 24,632,211-25,033,464 in AX_14 for 99 canine samples. Sites homozygous for the reference (0/0; blue) and alternate alleles (1/1; orange) are indicated along with heterozygous sites (0/1; white). Each column represents a single SNP (position indicated along top), while each row is a sample. Canid groupings are on the right of the matrix.

#### Refined identification of differentiated loci using demographic models and ancient genomes

Our results indicate that the use of village dogs in selection scans identifies novel candidate domestication loci. However, we further refined these F_ST_-based domestication scans to account for several shortcomings. First, rather than setting an empirical threshold at a ZF_ST_ score of 5, we created a neutral null model that captures key aspects of dog and wolf demographic history (Additional File 2: Figure S2; Additional File 3: Table S2; [34,40]). We identified 443 autosomal sliding windows with F_ST_ values that exceed the 99^th^ percentile of the neutral simulations (F_ST_ = 0.308; Additional File 2: Figure S3a; Additional File 2: Figure S4a). Second, reasoning that a true domestication sweep will be largely fixed among extant dogs with no recent wolf admixture, we calculated pooled heterozygosity (H_P_) in village dogs within the same window boundaries, and retained windows with a H_P_ lower than the 0.1^th^ percentile observed in our simulations (Additional File 2: Figure S4b). This heterozygosity filter removed 199 of the 443 windows (Additional File 2: Figure S3b). Finally, we excluded regions where the putatively selected haplotype is not found in ancient dog samples. To do this, we calculated the difference in dog H_P_ (ΔH_P_) with and without the inclusion of two ancient dog samples HXH, a 7ky old dog from Herxheim, Germany ([34]) and NGD, a 5ky old dog from Newgrange, Ireland ([21,34]); see Methods). Windows with ΔH_P_ greater than the 5^th^ percentile of all windows genome-wide (ΔH_P_ = −0.0036) were removed (Additional File 2: Figure S3c; Additional File 2: Figure S4c and d). Remaining overlapping windows were merged, resulting in 58 autosomal CDRs that encompass 18.65 Mbp of the genome and are within 50kb of 248 Ensembl gene models (Additional File 4: Table S3; Figure 3b).

#### Assessment of previously identified CDRs

We applied the same filtration parameters to putative dog domestication sweeps identified on autosomes in Axelsson et al. (N = 30; [26]) and Cagan and Blass (N = 5; [27]) (Additional File 2: Figure S5a and b). Since window coordinates of these studies may not precisely match our own, we selected the maximum F_ST_ value per locus from our village dog and wolf data. We then removed any locus with F_ST_, H_P_, and ΔH_P_ levels not passing our thresholds. Following these three filtration steps only 14 Axelsson, and 4 Cagan and Blass loci remained. Though few autosomal loci were reported in Cagan and Blass, 4 of the 5 loci passed and all have a corresponding sweep in either Axelsson or this study. These comparisons reinforce that sampling only or mostly breed dogs identifies many candidate sweeps that may have arisen post-domestication or as a result of breed formation. We also assessed the 349 loci identified by [29] by various statistics, and found that only 41 loci remained after similar filtrations (Additional File 2: Figure S5c)

### Increased resolution for domestication scans using the XP-CLR statistic

We further refined our search for domestication sweeps in village dogs using XP-CLR, a statistic developed to identify loci under selection based on patterns of correlated allele frequency differences between two populations [41]. Since we are searching for regions under selection in the dog genome, wolves were set as our reference population and XP-CLR was run on both simulated and real SNP datasets with a spacing of 2 kb, and a window size of 50 kb. Average XP-CLR values were calculated within 25 kb sliding windows (10 kb step size) for both datasets, and we retained 889 windows with scores greater than the 99th percentile from simulations (XP-CLR = 19.78; Additional File 2: Figure S6a). Using methods similar to those employed for F_ST_ scans, windows with village dog H_P_ values less than the 0.1^st^ simulation percentile (H_P_ = 0.0598) or where the ancient dog samples carried a different haplotype (ΔH_P_ filtration threshold at 5^th^ percentile = −0.0066) were eliminated (Additional File 2: Figure S6b-d; Additional File 2: Figure S3c). This resulted in 598 autosomal windows which we merged into 246 candidate loci, encompassing 10.81 Mb of genomic sequence (Figure 3a; Additional File 5: Table S4). These windows are located within 50 kb of 178 Ensembl gene models, and no SNPs with high F_ST_ within the intervals had deleterious effects on coding sequence, based on Variant Effect Predictor (Additional File 6: Table S5; [42]). Finally, the vast majority of the filtered XP-CLR sweeps (204/246) are novel to this study, 21 of which intersect F_ST_ loci from the non-outlier approach.

**Figure 3.**
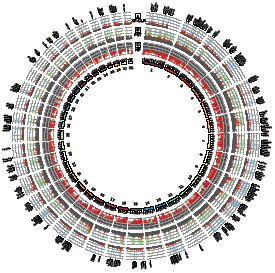
Circos plot of genome-wide selection statistics. Statistics from multiple selection scans are provided across the autosomes (chromosomes identifiers are indicated in the inner circle). (A) Averaged XP-CLR scores in 25kb windows across the genome. Windows with significant scores (greater than 99th percentile from simulations) are in red, and those that passed filtration are in blue. Genes within significant windows are listed above each region. (B) F_ST_ values calculated in 100kb windows. Values greater than the 99th percentile of simulations are in red. Windows that passed filtration are in green. (C) Averaged pooled heterozygosity (H_P_) scores in 25kb windows are indicated. Significant values (less than the 0.1th simulation percentile) are in red.

### Gene enrichment among 246 refined candidate domestication regions

We sought to identify gene sets and pathways enriched within our candidate domestication loci. Because of the increased resolution provided by XP-CLR (*i.e.* smaller window size), we focused the following gene analyses on the autosomal XP-CLR loci. Of these regions, 78% were within 50 kb of at least one gene. Based on 1,000 randomized permutations (see Methods), we found that the XP-CLR sweeps are not more likely to localize near genes than expected (p = 0.07), though the loci are near a greater total number of genes than random permutations (p = 0.003; Additional File 2: Figure S7a and b). Finally, gene set or pathway enrichment results can be influenced by gene size, as small genes are less likely to intersect with a sweep, potentially biasing toward enrichment of pathways that contain long genes. We observed that our sweeps contain genes of the similar average length as found in the randomized set (p > 0.05; Additional File 2: Figure S7c).

Gene ontologies were assigned to each dog Ensembl gene model using the BLAST2GO pipeline (see Methods). Enrichment tests were performed by the topGO R package [43] using the intersecting gene models as the test set, and the autosomal dog genes as the reference. In total, 292 significantly enriched GO categories were identified (p < 0.05) using the Parent-Child model of Fisher’s Exact Test [44]. To eliminate false positives, we also performed GO assignments and topGO enrichment tests for genes intersecting the randomized permutations and eliminated any XP-CLR GO category that was observed in more than 50 permuted enrichments (rate > 0.05) was excluded. After this filtration, 243 GO categories remained (Additional File 7: Table S6), 17 consisting of twenty or more XP-CLR loci. Lessening bias from singletons, we focused on the 174 GO terms represented by more than one XP-CLR locus. The top categories include GO categories associated with DNA replication (post-replication repair), transcriptional activity (STAT cascade, transcriptionally active chromatin), cellular localization, signaling, insulin secretion, and organismal development (midbody, post-embryonic animal organ morphogenesis). Additional biological pathways are highlighted below.

#### Candidate Genes Influencing Retinoic Acid Signaling

Retinoic acid (RA) is a signaling molecule that has numerous critical roles in development at the embryonic level, continuing into adult stages with roles including maintaining stem cell proliferation, tissue regeneration, and regulation of circadian rhythm [45,46]. The highest scoring XP-CLR locus centers upon *RAI1* (*Retinoic Acid-Induced 1;* Figure 4), a gene with numerous developmental functions in the RA pathway and the gene responsible for Smith-Magenis and Potocki-Lupski Syndromes in humans ([47,48]). Emphasizing the possible role of altered retinoic acid signaling in dog domestication, regulation of RA receptors were significantly enriched (GO:0048385; p = 0.0496). This category includes *NR2C1* (*Nuclear Receptor Subfamily 2 Group C Member 1*), essential for the development of early retina cells through regulation of early transcription factors that govern retinal progenitor cells such as RA receptors [49]. The second gene encodes Calreticulin, a protein involved in inhibition of both androgen and RA transcriptional activities [49,50]. Additionally, *Ncor2* (*Nuclear Receptor Corepressor 2*), found in In XP-CLR_209, increases cell sensitivity to RA when knocked out in mice [51].

**Figure 4.**
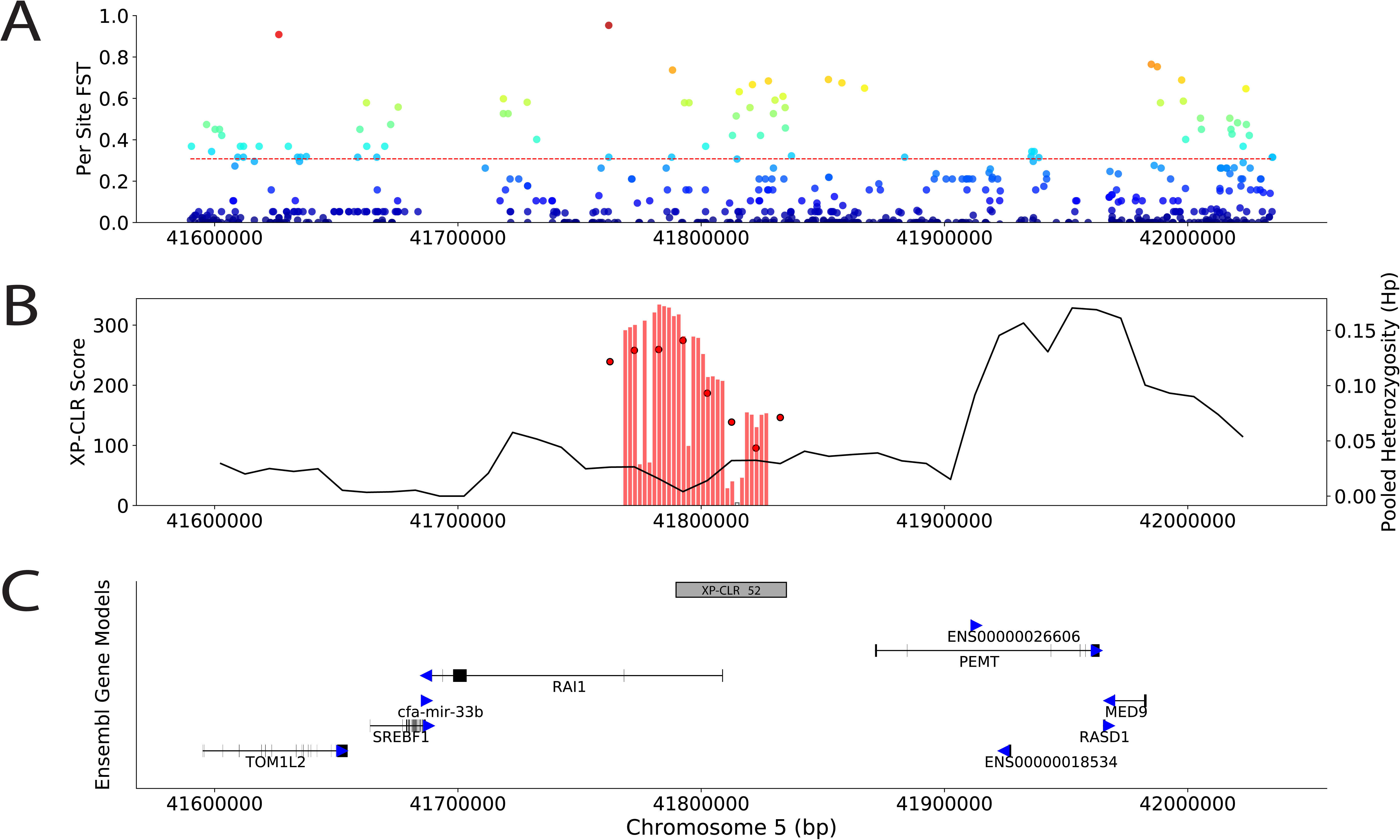
Selection scan statistics at the *RAI1* Locus. Selection scan statistics surrounding the *Retinoic Acid-Induced 1* (*RAI1*) locus (chr5: ~41.6-41.2 Mb) (A) Per site F_ST_ scores for all SNPs are indicated along with the F_ST_ significance threshold determined by the 99^th^ percentile of simulations (red dashed line). (B) Bars represent average XP-CLR scores in 25kb windows, those filled in red are less than the 0.1^th^ percentile from simulations. The black line indicates the average pooled heterozygosity (H_P_) values for the same window boundaries. (C) The significant XP-CLR locus (gray box) is presented relative to Ensembl gene models (black). Direction of each gene is indicated with blue arrows.

#### Candidate Genes Regulating Brain Development and Behavior

An abundance of genes and ontologies associated with neurological function, neurotransmission, and behavior were observed in our XP-CLR loci. GO categories such as neurotrophin binding, regulation of neurotransmitter levels, presynapse, as well as spines of both the neuron and dendrite were significantly enriched. Additionally, twelve genes, each from distinct XP-CLR loci, are linked to presynapse function (GO:0098793; p = 0.0143). Within these and additional loci, we observe numerous genes associated with neurotransmission. Representing two behavior-associated neurotransmitter pathways, the serotonin transporter *SLC6A4* and dopamine signaling members *GNAQ* and *ADCY6*(GO:0007212; p = 0.044) are also present. Genes associated with glutamate, the excitatory neurotransmitter, include *DGKI* (*Diacylglycerol kinase iota*; ranked 6th by XP-CLR), which regulates presynaptic release in glutamate receptors [52], and *GRIK3* (*Glutamate receptor, ionotropic, kainate 3*), a glutamate receptor [53]. Finally, *UNC13B* (*Unc-13 Homolog B*) is essential for competence of glutamatergic synaptic vesicles [54], and *CACNA1A* (*Calcium Voltage-Gated Channel Subunit Alpha1 A*) influences glutamatergic synaptic transmission [55].

In contrast to glutamate, GABA is the nervous system’s inhibitory neurotransmitter, and has been linked to the response to and memory of fear [53,56,57]. Genes in our XP-CLR loci relating to GABA include one of the two mammalian GABA biosynthetic enzymes *GAD2* (or *GAD65*; ranked 20th), the GABA receptor *GABRA4*, auxiliary subunit of GABA-B receptors *KCTD12* ([52,58]), and the GABA inhibitor osteocalcin (or *BGLAP*; [59]). Lastly, *T-Cell Leukemia Homeobox 3* (*TLX3*) is a key switch between glutamatergic and GABAergic cell fates [60].

#### Candidate Genes with Core DNA and RNA Functions

Pathways involved in core DNA and RNA processes are also well represented within the XP-CLR regions. Highly significant GO categories relate to the STAT (signal transducers and activators of transcription) cascade and its regulation (3^rd^, 6^th^, and 7^th^ top categories), represented by ten genes from nine unique XP-CLR loci including Interleukins (*IL22, IL26*), an Interleukin receptor (*IL31RA*), *SOCS3* (*Suppressor Of Cytokine Signaling 3*), and *CYP1B1* (*Cytochrome P450 1B1*). In addition to the STAT transcriptional activation pathway, genes involved in maintaining transcriptionally active chromatin, positively regulating transcription from RNA polymerase III promoter, and cryptic unstable transcript catabolism (CUT) are also enriched in our GO terms.

The GO category of RNA binding contains 59 genes from 52 unique loci and is highly enriched (p=0.014). This category contains genes involved in splicing of transcripts by both the major and minor splicing pathways. The eighth highest XP-CLR region harbors the gene *RNPC3* (*RNA Binding Region (RNP1, RRM) Containing 3*), the 65 KDa subunit of the U12 minor spliceosome, which is located ~55 kb downstream of *AMY2B* (Figure 5). Another core subunit, *SF3B1* (*Splicing Factor 3b Subunit 1*), belongs to both the minor and major (U2) spliceosome. Additional XP-CLR genes related to splicing and/or spliceosome function include *FRG1* [61], *DDX23* (alias *PRP28*; [62]), *CELF1* [63], *NSRP1*(alias *NSrp70*; [64,65]), and *SRSF11* (alias *P54*; [66]).

**Figure 5.**
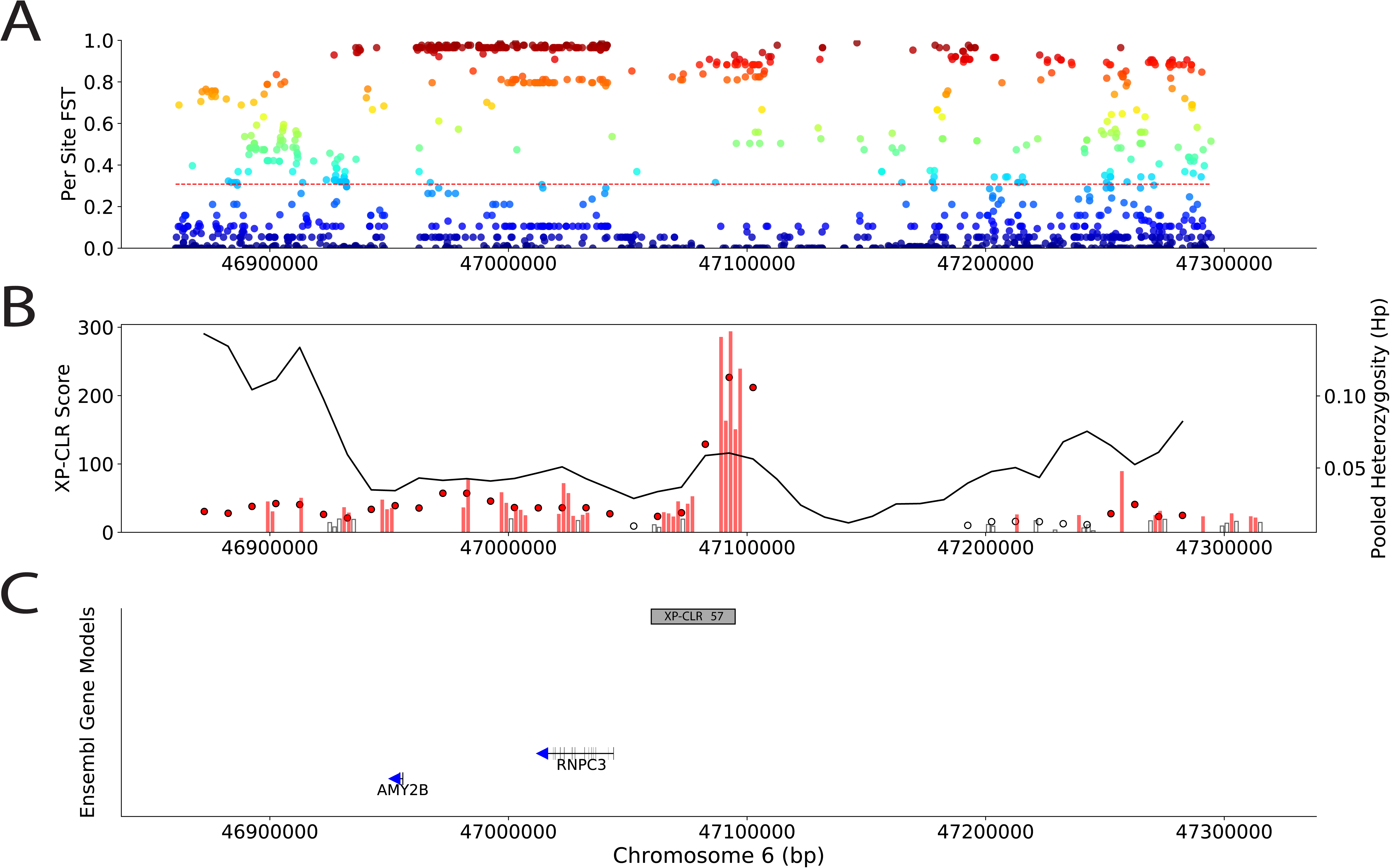
Selection scan statistics at the *RNPC3* Locus. Selection scan statistics surrounding the *RNA Binding Region (RNP1, RRM) Containing 3* (*RNPC3*) locus (chr5: ~46.9-47.3 Mb) (A) Per site F_ST_ scores for all SNPs are indicated along with the F_ST_ significance threshold determined by the 99^th^ percentile of simulations (red dashed line). (B) Bars represent average XP-CLR scores in 25kb windows, those filled in red are less than the 0.1^th^ percentile from simulations. The black line indicates the average pooled heterozygosity (H_P_) values for the same window boundaries. (C) The significant XP-CLR locus (gray box) is presented relative to Ensembl gene models (black). Direction of each gene is indicated with blue arrows.

#### Candidate Genes Involved In Cell Cycle Control

Nine genes involved in the regulation of cell cycle progression were located in XP-CLR loci (GO:1901990; p = 0.039), including *BIRC5* (or *survivin*) which is an apoptosis inhibitor [67,68]. Other enriched categories included postreplication repair (GO:0006301; p=0.013) and cell maturation (GO:0048469; p = 0.01299). Perturbations in any of these pathways could influence numerous biological functions such as cell proliferation, cell fate/death, and repair.

### Survey of copy-number variation between dogs and wolves

Copy-number variants have also been associated with population-specific selection and domestication in a number of species [5,69,70]. Since regions showing extensive copy-number variation may not be uniquely localized in the genome reference and may have a deficit of SNPs passing our coverage thresholds, we directly estimated copy number along the reference assembly and searched for regions of extreme copy number differences (see Methods). Using V_ST_, a statistic analogous to F_ST_ [69], we identified 67 regions of extreme copy number difference between village dogs and wolves which are within 50 kb of 89 unique genes (“Variant Candidate Domestication Regions” or VCDRs; Additional File 8: Table S7; see Additional File 9: Note 3). There was no overlap of VCDRs with regions identified through F_ST_ or XP-CLR.

Relative to randomly permuted intervals, the 67 VCDRs are more likely to be near genes (p < 0.01; Additional File 2: Figure S8a) but do not encompass more total genes than expected (p > 0.05; Additional File 2: Figure S8b). With methods implemented for XP-CLR loci, gene ontology enrichments and permutations were performed for genes within 50 kb of the VCDRs. GO enrichment analysis identified 111 significantly enriched categories (p < 0.05 using Parent-Child statistics; [43]). However, only 104 categories remained after filtration of GO categories found in >5% of random permutations (Additional File 10: Table S8). Five enriched terms were supported by more than two VCDRs including regulation of circadian rhythm (N=2 VCDRs), lipid droplet (N=3), calcium and calmodulin responsive adenylate (N=2), cyclase activity (N=2), steroid binding (N=2) and beta-catenin binding (N=2).

### Refined analysis of a large-scale structural variant located at the AMY2B locus

The top VCDR encompasses the *AMY2B gene*, which at increased copy number confers greater starch metabolism efficiency due to higher pancreatic amylase enzyme levels [5,38]. Quantitative PCR results have suggested an ancient origin for the *AMY2B* copy number expansion, as 7ky old Romanian dogs exhibit elevated *AMY2B* copy number [26,37]. However, read-depth analysis shows that the *AMY2B* tandem expansion is absent in 5-7ky old ancient European dogs [34]. However, read-depth profiles identified two large duplications, one of 1.9 Mb and the other of 2.0 Mb, that encompass *AMY2B* (Additional File 2: Figure S9). We quantified copy number at *AMY2B* itself and regions which discriminate the two segmental duplications in 90 dogs using digital droplet PCR (ddPCR). Copy number estimated through read depth strongly correlated with estimates from ddPCR (Additional File 2: Figure S10) confirming the presence of standing copy number variation of *AMY2B* in dogs (range of 2*n_AMY2B_* = 2-18) and distinguishing the two large-scale duplications (Additional File 2: Figure S11). The extreme *AMY2B* copy number expansion appears to be independent of the large-scale duplications, as ddPCR results show that some dogs without the large duplications still have very high *AMY2B* copy number (Additional File 2: Figure S11). Read-depth patterns at the duplication breakpoints indicated that NGD, the ancient Irish dog, harbored the 2.0 Mb duplication resulting in increased *AMY2B* copy number.

## Discussion

Genetic and archaeological data indicate that the dog was first domesticated from Eurasian gray wolves well over 10 kya [25,34,40,71]. Evidence suggests that the domestication process was complex and may have spanned thousands of years [3,71]. Through multiple analyses, we have identified regions that are strongly differentiated between modern village dogs and wolves and which may represent targets of selection during domestication. Our approach differs from previous studies in several ways including the use of village dogs rather than breed dogs, using neutral simulations to set statistical cutoffs, and filtering candidate loci based on ancient dog DNA data. Most (83%) of the 246 candidate domestication regions we identified are novel to our study, which we largely attribute to reduced signals associated with post-domestication breed formation. We argue that swept haplotypes identified in modern village dogs and also present in Neolithic dogs more likely represent signals of ancient selection events.

While the use of neutral simulations accounts for genetic diversity in both wild and domestic sampled populations, and better controls false positive rates than arbitrary empirical thresholds [29,72], several limitations are still apparent in our approach. The demographic model we used does not capture all aspects of dog history, does not include the X chromosome, and does not fit all aspects of the observed data equally well. This likely represents unaccounted for features of the data, such as unmodeled population structure, as well as technical issues such as reduced ascertainment of low frequency alleles due to sequencing depth. Although the inclusion of ancient samples allows for the removal of candidate sweeps that are unique to modern dogs, this approach is limited by the narrow temporal (5-7 kya) and geographic (restricted to Europe) sampling offered by the available data. Even though most selected alleles likely preexisted in the ancestral wolf population, our approach identifies regions where modern dogs all share the same haplotype. However, even when selection acts on preexisting mutation a single haplotype often reaches fixation [73], consistent with the variation patterns we identify across village dog populations.

Analysis of genes near the 246 loci identified using XP-CLR identified numerous overrepresented ontologies including developmental, metabolic, signaling, and neurological pathways (Additional File 7: Table S6). Importantly, our gene annotations were obtained directly through established BLAST2GO pipelines [74], rather than assignment of function via homologous human genes, as has been employed in earlier dog domestication studies [26,28,29]. We additionally performed permutations to remove functional categories likely to be observed by chance. Importantly, the genomic positions of the genes in most enriched categories are not co-localized, implying that the observed enrichment is not an artifact of the clustering of genes with similar functions.

### The role of the neural crest in dog domestication

Our CDRs include 52 genes also identified in analysis of other domesticated or self-domesticated animals [9,11,17,75–79], including four genes (*RNPC3, CUEDC1, GBA2, NPR2*) in our top 20 XP-CLR loci. No gene was found in more than three species, consistent with the hypothesis that no single domestication gene exists [18]. Although the overlap of specific genes across species is modest, there are many enriched gene pathways and ontologies shared in domesticates including neurological and nervous system development, behavior, reproduction, metabolism, and pigmentation [10,11,17,7,5,80,81]. We attribute these patterns to the domestication syndrome (DS), a phenomenon where diverse traits, manifested in vastly different anatomical zones, appear seemingly disconnected, yet are maintained across domesticates. Two possible modes of action could generate the DS phenotypes while still displaying the genome-wide distribution of sweeps. The first would require independent selection events for distinct traits at numerous loci. Alternatively, selection could have acted on considerably fewer genes that are members of early-acting developmental pathways with broad phenotypic effects.

For these reasons, the role of the neural crest in animal domestication has gained support from researchers over recent years [18,82]. In 2014, Wilkins et al. [18]established that the vast array of phenotypes displayed in the animal DS mirror those exhibited in mild human neurocristopathies, whose pathology stems from aberrant differentiation, division, survival, and altered migration of neural crest cells (NCCs). These cells are multipotent, transient, embryonic stem cells that are initially located at the crest (or dorsal border) of the neural tube. The initiation and regulation of neural crest development is a multi-stage process requiring the actions of many early-expressed genes including the Fibroblast Growth Factor (*Fgf*), Bone Morphogenic Protein (*Bmp*), Wingless (*Wnt*), and *Zic* gene families [83]. Several of the genes identified in our analysis are involved in this transition including members of the Fgf (*Fgf1*) family as well as a transcription factor (*TCF4*; [84]), inhibitors (RRM2; *NPHP3*; [85,86]) and regulators (*LGR5; [87]*) of the Wnt signaling pathways.

Following induction, NCCs migrate along defined pathways to various sites in the developing embryo. Assignment of identity and the determination of migration routes relies on positional information provided by external signaling cues [88,89]. *KCTD12, CLIC4, PAK1, NCOR2, DOCK2*, and *EXOC7* are examples of such genes that are linked to the determination of symmetry, polarity, and/or axis specification [90–94]. Together, our results suggest that early selection may have acted on genes essential to the initiation of the neural crest and the definition of migration routes for NCCs.

#### NCC-Derived Tissues Linked to Domestication Syndrome Phenotypes

Once in their final destinations, NCC further differentiate as the precursors to many tissues in the developing embryo. Most of the head, for example, arises from NCCs including craniofacial bones, cartilage, and teeth [95,96]. Ancient dog remains indicate that body size, snout lengths, and cranial proportions of dogs considerably decreased compared to the wolf ancestral state following early domestication [97]. Further, these remains indicate jaw size reduction also occurred, as evidenced by tooth crowding [97]. Such alterations are consistent with the DS and implicate aberrant NCC migration since decreases in the number of NCCs in facial primordia are directly correlated with reductions in mid-face and jaw sizes [18,98]. Genes associated with both craniofacial and tooth development in vertebrates are found in our candidate loci including *SCUBE1*, which is essential in craniofacial development of mice, and *SATB2*, which has roles in patterning of the developing branchial arches, palate fusion, and regulation of *HOXa2* in the developing neural crest [99–101]. Lastly, when knocked out in mice, *Bicoid*-related homeodomain factor *PITX1* not only affected hindlimb growth, but also displayed craniofacial abnormalities such as cleft palate and branchial arch defects [102] and influences vertebrate tooth development [103].

Insufficient cartilage, a NCC-derived tissue [96] that consists of chondrocytes and collagen, in the outer ear of humans results in a drooping ear phenotype [104] linked to numerous NC-associated neurocristopathies (e.g. Treacher Collins, Mowat-Wilson, etc.). Analogously, compared to the pricked ears of wolves, dogs predominantly have “floppy” ears [105], a hallmark feature of domesticates [18]. *SERPINH1*, a collagen-binding protein, is embryonically lethal when ablated in mice [106], and appears to be required for chrondrocyte maturation [107]. Alterations of activity by genes such as *SERPINH1* and those regulating NCC migration may have reduced the numbers of NCCs in dog ears, contributing to the floppy phenotype [18].

#### Mis-regulation of gene expression may contribute to DS phenotypes

Similar to other domestication scans [6,9,19], we did not find SNPs deleteriously altering protein sequence in our predicted sweeps, indicating that gene loss did not have a significant role in dog domestication. Instead, we hypothesize that alterations in gene regulatory pathways or the regulation of transcriptional activity could contribute to broad DS phenotypes. In addition to over-representations of genes influencing transcriptional control (STAT cascade) and maintenance of transcriptionally active chromatin, we identified two components of the minor spliceosome; *RNPC3* and *Sf3b1. RNPC3*, which affects early development and is linked to dwarfism (Isolated Growth Hormone Deficiency; [108]), is also under selection in cats and humans [17,77]. Absence of *Sf3b1* disrupts proper NCC specification, survival and migration [109]. A further example of the role of splicing in NC development is that mutations in *U4atac*, a U12 snRNA subunit gene missing in the current dog annotation, causes Taybi-Lindner syndrome (TALS) in humans. Phenotypes of this syndrome resemble those of the DS including craniofacial, brain, and skeletal abnormalities [110]. Thus, proper splicing, particularly for transcripts processed by the minor spliceosome, is required for proper NC function and development.

#### Genes associated with neurological signaling, circadian rhythms, and behavior

Tameness or reduced fear toward humans was likely the earliest trait selected for by humans during domestication [3,111,112]. Recapitulating such selection, numerous physiological and morphological characteristics, including DS phenotypes (*i.e.* floppy ears, altered craniofacial proportions, and unseasonal timing for mating), appeared within twenty generations when researchers selected only for tameness in a silver fox breeding population [1,113]. As the progenitors for the adrenal medulla, which produces hormones associated with the “fight-or-flight” response, hypofunction of NCCs can lead to changes in the tameness of animals [18]. The link between tameness and the NC suggests that changes in neural crest development could have arisen first, either through direct selection by humans for desired behaviors or via the “self-domestication” [114,115] of wolves that were more docile around humans.

Numerous candidate genes influencing neurological function and behavioral responses were enriched in XP-CLR loci including genes in the dopamine, serotonin, glutamate, and GABA neurotransmission pathways. These, coupled with enriched GO categories relating to neuronal projections (*e.g.* neuron and dendrite spines), suggest that neuronal signaling may have been differentially impacted in dogs through domestication, in agreement with Li et al. [116].

Our top XP-CLR locus was located within *RAI1*, the gene for Smith-Magenis syndrome (SMS; [117]), a disorder that is associated with craniofacial and skeletal deformations, aggression, developmental delays, altered circadian rhythms, and intellectual disabilities [118]. A recent study of breed dogs and wolves by vonHoldt et al. [119] linked structural variants near the *WBSCR17* gene to behavioral changes in dogs. *WBSCR17* is associated with another neurodevelopmental disorder, Williams-Beuren syndrome, that displays hypersociability in humans [120]. Williams-Beuren and Smith-Magenis syndromes each display multiple features associated with improper NCC development [118,121]. In *Xenopus*, disruption of the transcription factors *RAI1* and *WSTF* (*Williams-Beuren Syndrome Transcription Factor*, a gene also disrupted in Williams-Beuren syndrome) negatively impacts proper NCC migration, recapitulating the craniofacial defects associated with the syndromes [122,123]. Furthermore, both syndromes involve perturbations in sleep patterns [124–127], and *RAI1* has been shown to regulate the circadian rhythm pathway [126]. Alterations of sleep patterns occurred in dogs as they shifted from the ancestral nocturnal state of wolves, to that of the diurnal lifestyle also exhibited by humans. In domesticated silver foxes selected for tameness, levels of circadian rhythm determinants (*e.g.* melatonin and serotonin) were significantly altered compared to wild foxes [128–130]. This suggests possible overlap between genes influencing behavior, circadian rhythms, and the Domestication Syndrome.

## Conclusions

By comparing village dogs and wolves, we identified 246 candidate domestication regions in the dog genome. Analysis of gene annotations in these regions suggests that perturbation of crucial neural crest signaling pathways results in the broad phenotypes associated with the Domestication Syndrome. Additionally, these findings link transcriptional regulation and splicing to alterations in cell differentiation, migration, and neural crest development. Altogether, we conclude that while primary selection during domestication likely targeted tameness, genes that contribute to determination of this behavioral change are also involved in critical, far-reaching pathways that conferred drastic phenotypic changes in dogs relative to their wild counterparts.

## Methods

### Sample Processing and Population Structure Analysis

Genomes of canids (Additional File 11: Table S9) were processed using the pipeline outlined in [34] to produce a data set of single nucleotide polymorphisms (SNPs) using GATK [131]. Thirty-seven breed dogs, 45 village dogs, and 12 wolves were selected from the samples described in Botigue et al. [34], and ADMIXTURE [132] was utilized to estimate the levels of wolf-dog admixture within this subset. To account for LD, the data was thinned with PLINK v1.07 (indep-pairwise 50 10 0.1; [133]), where SNPs with an R^2^ value over 0.1 were removed in 50kb windows, sliding 10 sites at a time. The remaining 1,030,234 SNPs were used in five independent ADMIXTURE runs using different seeds, for up to five ancestral populations (K = 1-5). K = 3 had the lowest average cross validation error (0.0373) from the five runs, and was therefore the best fit for the data. To eliminate noise in subsequent analyses, we removed all village dogs with greater than 5% wolf ancestry and wolves with greater than 5% dog ancestry. Fifty-four samples remained following this filtration.

Following elimination of admixed samples, we called SNPs in 43 village dogs and 11 gray wolves using GATK (v. 3.4-46; [131]). Using the GATK VQSR procedure, we identified a high quality variant set such that 99% of positions on the Illumina canine HD array were retained. VQSR filtration was performed separately for the autosomes + chrX pseudoautosomal region (PAR) and the non-PAR region. SNPs within 5 bp of an indel identified by GATK were also removed. We further excluded sites with missing genotype calls in any sample, triallelic sites, and X-nonPAR positions where any male sample was called as heterozygous. The final SNP set contained 7,657,272 sites.

Using these SNPs, we removed samples that exhibited over 30% relatedness following identity by state (IBS) analysis with PLINK v1.90 (min 0.05; [133]). Only one sample (*mxb*), was removed from the sample set, a sample known to be related to another mexican wolf in the dataset. Principal component analyses were completed on the remaining 53 samples (43 dogs and 10 wolves) using smartpca, a component of Eigensoft package version 3.0 [134] after randomly thinning the total SNP set to 500,000 sites using PLINK v.1.90 [135]. Once PCA confirmed clear genetic distinctions between these dogs and wolves, this final sample set was used for subsequent analyses. The SNP set was further filtered for the selection scans to remove rare alleles (minor allele frequencies < 3 out of possible 106 alleles, or 0.028). Finally, village dog and wolf allele frequencies were calculated separately using VCFtools ([136].

### Demographic Model and Simulations

Simulations of dog and wolf demographic history were performed using msprime v.0.4.0 [137]. For each autosome, 75 independent simulations were performed using independent random seeds and a pedigree based genetic map [138]. A mutation rate of 4 x10-^9^ per site per generation with a generation time of 3 years was assumed. The 53 samples were modeled as coming from 10 lineages with population histories adapted from [34,40] (Additional File 2: Figure S2; Additional File 3: Table S2). The simulation is designed to capture key aspects impacting dog and wolf diversity, rather than a definitive depiction of their demography. Resulting simulated SNP sets were filtered for minor allele frequency and randomly thinned to have the same number of SNPs per chromosome as the real SNP datasets used in F_S_T, XP-CLR, and H_P_ calculations.

### F_ST_ Selection Scans

Dog and wolf allele counts generated above were used to calculate the fixation index (FST) using the Hudson estimator derived in [139]with the formula: F_S_T = ((□_1_ - □_2_) -(□1(1 - □_1_) / □1 - 1) - (□_2_(1 - □_2_) / □_2_ - 1)) / (□1(1 - □_2_) + □_2_(1 - C1)) where n_n_ is the allele frequency in population x, and □_□_ is the number of individuals in population x. With this equation, the X chromosome could be included in F_S_T calculations. A custom script calculated the per site F_S_T across the genome for both the real and 75 simulated SNP sets. Due to differences in effective population size and corresponding expected levels of genetic drift, analyses were performed separately for the X-non-PAR. Ratio of averages for the resulting F_S_T values were calculated in 200kb sliding windows with 50kb step sizes, and we required each window to contain at least 10 SNPs. Additionally, we calculated per site FST for each SNP that did not have missing data in any sample.

FST loci filtration was completed differently for the outlier and non-outlier approach. For the outlier F_S_T approach, the windows were Z-transformed and only windows with Z scores ≥ 5 standard deviations were deemed significant for autosomal and X-PAR loci, and ≥ 3 for the X-NonPAR. Significance thresholds for the non-outlier approach were determined as the 99th percentile from F_S_T score distributions from the simulated genomes. Overlapping windows passing these thresholds were merged.

### Pooled Heterozygosity (HP) and ΔHP Calculations

Per window, dog allele frequencies were used to calculate pooled heterozygosity (H_P_) using the following formula from [6]: 2Σ*n*_MAJI_Σ*n*_MIN_/(Σ*n*_MAJ_□+□Σ*n*_MIN_)^2^, where Σ*n*_MAJ_ is the sum of major and Σ*n*_MIN_ minor dog alleles, respectively, for all sites in the window. Significance thresholds for window filtration was set as the 0.1^th^ percentile of the H_P_ distribution from the simulated genomes. The change in H_P_ (or ΔH_P_) was calculated as the difference in ΔH_P_ with and without the inclusion of the two ancient dog samples (HXH and NGD). The 5 ky old German dog (CTC) was not included in this analysis due to known wolf admixture [34]. Windows with AH_P_ greater than the 5^th^ percentile observed genome wide were removed.

### XP-CLR Selection Scans

XP-CLR was calculated using dog and wolf allele frequencies at sites described above [41]. This analysis requires separate genotype files for each population, and a single SNP file with positions of each SNP and their genetic distance (in Morgans), which were determined through linear extrapolation from the pedigree-based recombination map from [138]. Wolves were set as the reference population, and XP-CLR was run on both the simulated and real SNP sets with a grid size of 2 kb, and a window sizes of 50 kb. Windows that did not return a value (failed) or did not have at least five grids were removed. Average XP-CLR scores were calculated in 25 kb windows (step size = 10kb). Filtration of real windows with averages less than the 99^th^ percentile of averaged simulation scores was performed. All remaining adjacent windows were merged if they were within 50 kb distance (*i.e.* one sliding window apart).

### Visualization of sweeps

Forty-six additional canines (e.g. dog breeds, jackals, coyotes; Additional File 11: Table S9) were genotyped at candidate loci identified in this study, as well as those from [26,27], [27], and [29] using autosomal SNPs previously called in [34]. SNPs within CDRs of interest were extracted from the SNP dataset using the PLINK make-bed tool with no missing data filter. Per sample, each SNP was classified as 0/0, 0/1, or 1/1 at all loci (1 representing the non-reference allele), and this genotype data was stored in Eigenstrat genotype files, which were generated per window using convertf (EIGENSOFT package; [140]). Custom scripts then converted the Eigenstrat genotype files into matrices for visualization using matrix2png [141].

### Gene Enrichment and Variant Annotation

Coordinates and annotations of dog gene models were obtained from the Ensembl release 81 (ftp.ensembl.org/pub/release-81/ and useast.ensembl.org/Canis_familiaris/Info/Index, respectively), and a non-redundant annotation set was determined. The sequence of each Ensembl protein was BLASTed against the NCBI non-redundant database (blastp outfmt 5 evalue 1e-3 word_size 3 show_gis max_hsps_per_subject 20 num_threads 5 max target_seqs 20) and all blastp outputs were processed through BLAST2GO [142] with the following parameters: minimum annotation cut-off of 55, GO weight equal to 5, BLASTp cut-off equal to 1e^−6^, HSP-hit cut-off of 0, and a hit filter equal to 55. Positions of all predicted domestication loci (from this and previous studies) were intersected with the coordinates of the annotated Ensembl canine gene set to isolate genes within the putatively swept regions. Gene enrichment analyses were performed on these gene sets using topGO [43], and we utilized p-values produced by the parent-child test [44]. The predicted effects of SNP variants were obtained by the processing of the total variant VCF file of all canine samples by Variant Effect Predictor (VEP; [42]).

### Copy Number Estimation Using QuicK-mer and fastCN

We implemented two CN estimation pipelines to assess copy-number in village dogs and wolves using the depth of sequencing reads. The first, fastCN, is a modified version of existing pipelines that considers multi-mapping reads to calculate CN within 3kb windows (Additional File 9: Note 1). By considering multi-mapping reads, copy-number profiles will be shared among related gene paralogs, making it difficult to identify specific sequences that are potentially variable. The second pipeline we employed, QuicK-mer, a map-free approach based on k-mer counting which can accurately assess CN in a paralog-sensitive manner (Additional File 9: Note 2). Both pipelines analyze sequencing read-depth within predefined windows, apply GC-correction and other normalizations, and are able to convert read depth to a copy-number estimate for each window. The signal-to-noise ratio (SNR), defined as the mean depth in autosomal control windows divided by the standard deviation, was calculated for each sample. The CN states called by both the QuicK-mer and fastCN pipelines were validated through comparison with aCGH data from [143]. Regions with copy number variation between samples in the aCGH or WGS data were selected for correlation analysis (Additional File 9: Note 3).

### V_ST_ Selection Scans

V_ST_ values [69] were calculated for genomic windows with evidence of copy number variation. VST values were Z-transformed and we identified outlier regions as windows exhibiting at least a 1.5 copy number range across all samples, and ZV_ST_ scores greater than 5 on the autosomes and the X-PAR, or greater than 3 in the X-nonPAR. Prior to analysis, estimated copy numbers for male samples on the non-PAR region of the X were doubled. Outlier regions spanning more than one window were then classified as variant candidate domestication regions (VCDRs) (Additional File 8: Table S7). A similar analysis was performed for the unplaced chromosomal contigs in the CanFam3.1 assembly (Additional File 12: Table S10). See Additional File 9: Note 3 for additional methods and details.

### Amylase Structural Variant Analysis

We estimated copy-number using short-read sequencing data from each canine listed in Additional File 11: Table S9. Copy number estimates for the *AMY2B* gene using fastCN were based on a single window located at chrUn_AAEX03020568: 4873-8379. See Supplementary Methods: Note 3.5.1 for further methods and results. Digital droplet PCR (ddPCR) primers were designed targeting overlapping 1.9Mb and 2.0Mb duplications, the *AMY2B* gene, and a copy number control region (chr18: 27,529,623-27,535,395) found to have a copy number of 2 in all sampled canines by QuicK-mer and fastCN. Copy number for each target was determined from ddPCR results from a single replication for 30 village dogs, 3 New Guinea singing dogs, and 5 breed dogs (Additional File 13: Table S11) and averaged from two replicates for 48 breed dogs (Additional File 14: Table S12). For more details on primer design, methods and results for the characterization of the *AMY2B* locus, see Additional File 9: Note 3.

## Acknowledgements

We thank Shiya Song for advice and assistance in the processing of canid variation data and Laura Botigue for discussion of results utilizing ancient DNA.

## Funding

This work was supported by National Institutes of Health (R01GM103961 to AB and JMK, and T32HG00040 AP). DNA samples and associated phenotypic data were provided by the Cornell Veterinary Biobank, a resource built with the support of NIH grant R24GM082910 and the Cornell University College of Veterinary Medicine.

## Availability of data and materials

The datasets supporting the conclusions of this article are available in the article and its additional files, as well as a custom UCSC track hub (raw.githubusercontent.com/KiddLab/Pendleton_2018_Selection_Scan/master/Selection_track_hub.txt). Software (fastCN and QuicK-mer) implemented in this article are available for download in a GitHub repository (github.com/KiddLab/). Pre-computed 30-mers from the dog, human, mouse, and chimpanzee genomes can be downloaded from kiddlabshare.umms.med.umich.edu/public-data/QuicK-mer/Ref/ for QuicK-mer processing. Genome sequence data for three New Guinea singing dogs was published under project ID SRP034749 in the Short Read Archive.

## Author contributions

JMK, ALP, and FS designed the study. JMK oversaw the study. Selection scans were performed by ALP, AT, and FS. AT and ALP assessed population structure. CNVs were estimated by FS and JMK. Functional annotations and enrichment analyses were performed by ALP. FS processed aCGH data. KV processed ancient dog samples. Samples and genome sequences were provided by KV, AB and JMK. SE performed the DNA extractions, library generation, and ddPCR analyses. ALP, FS, and JMK wrote the paper with input from the other authors.

## Competing interests

ARB is a cofounder and officer of Embark Veterinary, Inc., a canine genetics testing company.

## Abbreviations

aCGH: array comparative genomic hybridization
CDR: candidate domestication region
chrUn: chromosome unknown
CN: copy number
CNV: copy number variation
ddPCR: droplet digital polymerase chain reaction
DS: Domestication Syndrome
GO: gene ontology
H_P_: pooled heterozygosity
NC: neural crest
NCC: neural crest cell
qPCR: quantitative polymerase chain reaction
SNP: single-nucleotide polymorphism
SNR: signal to noise ratio
VCDR: V_ST_ candidate domestication region
XP-CLR: cross-population composite likelihood ratio.

## Supplementary Material

**Additional File 1:** Table S1 - Coordinates and annotations of F_ST_ loci identified through outlier approach

**Additional File 2:** Supplemental figures S1-S11

**Additional File 3:** Table S2 - Parameters incorporated into demographic model for neutral evolution in village dog and wolf populations

**Additional File 4:** Table S3 - Coordinates and annotations of outlier F_ST_ loci identified through simulation approach

**Additional File 5:** Table S4 - Coordinates and annotations of XP-CLR candidate domestication regions identified through simulation approach

**Additional File 6:** Table S5 - Predicted SNP effects (per Variant Effect Predictor) for sites in XP-CLR candidate domestication regions

**Additional File 7:** Table S6 - Gene enrichment results for XP-CLR candidate domestication regions

**Additional File 8:** Table S7 - Coordinates of V_ST_ candidate domestication regions (VCDRs)

**Additional File 9:** Notes 1-3 providing supplementary methods and results of copy number analysis

**Additional File 10:** Table S8 - Gene enrichment results for V_ST_ candidate domestication regions (VCDRs)

**Additional File 11:** Table S9 - Description and accession numbers for canine genomes processed in this study

**Additional File 12:** Table S10 - Coordinates of V_ST_ candidate domestication regions on chromosome unknown

**Additional File 13:** Table S11 - ddPCR results from 30 village, 3 New Guinea singing, and 5 breed dog samples of AMY2B segmental duplications

**Additional File 14:** Table S12 - ddPCR results from 48 breed dog samples of AMY2B segmental duplications

**Additional File 15:** Plots displaying aCGH probe intensity correlations with in silico copy number estimates.

**Additional File 16:** Supplemental QuicK-mer validation figures.

## Supplementary Figures

**Figure S1**-Z transformed F_ST_ scores for Cagan and Blass locus.

**Figure S2**-Demographic model for village dog and wolf populations used in neutral simulations

**Figure S3**-Filtration pipeline implemented for F_ST_ and XP-CLR windows

**Figure S4**-Distribution of selection scan statistics for real and simulated F_ST_ windows

**Figure S5**-Filtration pipeline implemented for Axelsson, Cagan and Blass, and Freedman CDRs

**Figure S6**-Distribution of selection scan statistics for real and simulated XP-CLR windows

**Figure S7**-Gene intersect statistics from randomized permutations of XP-CLR gene positions

**Figure S8**-Gene intersect statistics from randomized permutations of V_ST_ gene positions

**Figure S9**-Read-depth profiles at the AMY2B locus highlights large-scale structuralvariant

**Figure S10**-Correlations between the ddPCR and read-depth estimated copy-number for AMY2B and associated segmental duplications

**Figure S11**-results for the AMY2B gene, 1.9 Mb duplication, and the 2.0 Mb duplication

## References

1. Trut L. Early Canid Domestication: The Farm-Fox Experiment. Am. Sci. 1999;87:160.

2. Germonpré M, Sablin MV, Lázničková-Galetová M, Després V, Stevens RE, Stiller M, et al. Palaeolithic dogs and Pleistocene wolves revisited: a reply to Morey (2014). J. Archaeol. Sci. 2015;54:210–6.

3. Larson G, Burger J. A population genetics view of animal domestication. Trends Genet. 2013;29:197–197.

4. Harlan JR. Crops & Man 7 Jack R. Harlan. 1975.

5. Axelsson E, Ratnakumar A, Arendt M-L, Maqbool K, Webster MT, Perloski M, et al. The genomic signature of dog domestication reveals adaptation to a starch-rich diet. Nature. 2013;495:360–360.

6. Rubin C-J, Zody MC, Eriksson J, Meadows JRS, Sherwood E, Webster MT, et al. Whole-genome resequencing reveals loci under selection during chicken domestication. Nature. 2010;464:587–587.

7. Li M, Tian S, Jin L, Zhou G, Li Y, Zhang Y, et al. Genomic analyses identify distinct patterns of selection in domesticated pigs and Tibetan wild boars. Nat. Genet. 2013;45:1431–1431.

8. Cagan A, Blass T. Identification of genomic variants putatively targeted by selection during dog domestication. BMC Evol. Biol. 2016;16:10.

9. Rubin C-J, Megens H-J, Martinez Barrio A, Maqbool K, Sayyab S, Schwochow D, et al. Strong signatures of selection in the domestic pig genome. Proc. Natl. Acad. Sci. U. S. A. 2012;109:19529–19529.

10. Qiu Q, Wang L, Wang K, Yang Y, Ma T, Wang Z, et al. Yak whole-genome resequencing reveals domestication signatures and prehistoric population expansions. Nat. Commun. 2015;6:10283.

11. Fariello M-I, Servin B, Tosser-Klopp G, Rupp R, Moreno C, International Sheep Genomics Consortium, et al. Selection signatures in worldwide sheep populations. PLoS One. 2014;9:e103813.

12. Fang M, Larson G, Ribeiro HS, Li N, Andersson L. Contrasting mode of evolution at a coat color locus in wild and domestic pigs. PLoS Genet. 2009;5:e1000341.

13. Wang Z, Yonezawa T, Liu B, Ma T, Shen X, Su J, et al. Domestication relaxed selective constraints on the yak mitochondrial genome. Mol. Biol. Evol. 2011;28:1553–1553.

14. Cheng T, Fu B, Wu Y, Long R, Liu C, Xia Q. Transcriptome sequencing and positive selected genes analysis of Bombyx mandarina. PLoS One. 2015;10:e0122837.

15. Gray MM, Granka JM, Bustamante CD, Sutter NB, Boyko AR, Zhu L, et al. Linkage disequilibrium and demographic history of wild and domestic canids. Genetics. 2009;181:1493–505.

16. Amaral AJ, Megens H-J, Crooijmans RPMA, Heuven HCM, Groenen MAM. Linkage disequilibrium decay and haplotype block structure in the pig. Genetics. 2008;179:569–569.

17. Montague MJ, Li G, Gandolfi B, Khan R, Aken BL, Searle SMJ, et al. Comparative analysis of the domestic cat genome reveals genetic signatures underlying feline biology and domestication. Proc. Natl. Acad. Sci. U. S. A. 2014;111:17230–17230.

18. Wilkins AS, Wrangham RW, Fitch WT. The “domestication syndrome” in mammals: a unified explanation based on neural crest cell behavior and genetics. Genetics. 2014;197:795–795.

19. Carneiro M, Rubin C-J, Di Palma F, Albert FW, Alföldi J, Barrio AM, et al. Rabbit genome analysis reveals a polygenic basis for phenotypic change during domestication. Science. 2014;345:1074–1074.

20. Fallahsharoudi A, de Kock N, Johnsson M, Ubhayasekera SJKA, Bergquist J, Wright D, et al. Domestication Effects on Stress Induced Steroid Secretion and Adrenal Gene Expression in Chickens. Sci. Rep. 2015;5:15345.

21. Frantz LAF, Mullin VE, Pionnier-Capitan M, Lebrasseur O, Ollivier M, Perri A, et al. Genomic and archaeological evidence suggest a dual origin of domestic dogs. Science. 2016;352:1228–31.

22. Shannon LM, Boyko RH, Castelhano M, Corey E, Hayward JJ, McLean C, et al. Genetic structure in village dogs reveals a Central Asian domestication origin. Proc. Natl. Acad. Sci. U. S. A. 2015;112:13639–13639.

23. Wang G-D, Zhai W, Yang H-C, Wang L, Zhong L, Liu Y-H, et al. Out of southern East Asia: the natural history of domestic dogs across the world. Cell Res. 2016;26:21–21.

24. vonHoldt BM, Pollinger JP, Lohmueller KE, Eunjung H, Parker HG, Pascale Q, et al. Genome-wide SNP and haplotype analyses reveal a rich history underlying dog domestication. Nature. 2010;464:898–898.

25. Skoglund P, Ersmark E, Palkopoulou E, Dalén L. Ancient wolf genome reveals an early divergence of domestic dog ancestors and admixture into high-latitude breeds. Curr. Biol. 2015;25:1515–1515.

26. Axelsson E, Ratnakumar A, Arendt M-L, Maqbool K, Webster MT, Perloski M, et al. The genomic signature of dog domestication reveals adaptation to a starch-rich diet. Nature. 2013;495:360–360.

27. Cagan A, Blass T. Identification of genomic variants putatively targeted by selection during dog domestication. BMC Evol. Biol. 2016;16:10.

28. Wang G-D, Zhai W, Yang H-C, Fan R-X, Cao X, Zhong L, et al. The genomics of selection in dogs and the parallel evolution between dogs and humans. Nat. Commun. 2013;4:1860.

29. Freedman AH, Schweizer RM, Ortega-Del Vecchyo D, Han E, Davis BW, Gronau I, et al. Demographically-Based Evaluation of Genomic Regions under Selection in Domestic Dogs. PLoS Genet. 2016;12:e1005851.

30. Boyko AR. The domestic dog: man’s best friend in the genomic era. Genome Biol. 2011;12:216.

31. Pilot M, Malewski T, Moura AE, Grzybowski T, Oleriski K, Ruse A, et al. On the origin of mongrels: evolutionary history of free-breeding dogs in Eurasia. Proc. Biol. Sci. 2015;282:20152189.

32. Freedman AH, Gronau I, Schweizer RM, Ortega-Del Vecchyo D, Han E, Silva PM, et al. Genome sequencing highlights the dynamic early history of dogs. PLoS Genet. 2014;10:e1004016.

33. Marsden CD, Ortega-Del Vecchyo D, O’Brien DP, Taylor JF, Ramirez O, Vilà C, et al. Bottlenecks and selective sweeps during domestication have increased deleterious genetic variation in dogs. Proc. Natl. Acad. Sci. U. S. A. 2016;113:152–7.

34. Botigué LR, Song S, Scheu A, Gopalan S, Pendleton AL, Oetjens M, et al. Ancient European dog genomes reveal continuity since the Early Neolithic. Nat. Commun. 2017;8:16082.

35. Kelley JL, Madeoy J, Calhoun JC, Swanson W, Akey JM. Genomic signatures of positive selection in humans and the limits of outlier approaches. Genome Res. 2006;16:980–9.

36. Arendt M, Cairns KM, Ballard JWO, Savolainen P, Axelsson E. Diet adaptation in dog reflects spread of prehistoric agriculture. Heredity. 2016;117:301–6.

37. Ollivier M, Tresset A, Bastian F, Lagoutte L, Axelsson E, Arendt M-L, et al. Amy2B copy number variation reveals starch diet adaptations in ancient European dogs. R Soc Open Sci. 2016;3:160449.

38. Arendt M, Fall T, Lindblad-Toh K, Axelsson E. Amylase activity is associated with AMY2B copy numbers in dog: implications for dog domestication, diet and diabetes. Anim. Genet. 2014;45:716–22.

39. Alexander DH, Novembre J, Lange K. Fast model-based estimation of ancestry in unrelated individuals. Genome Res. 2009;19:1655–64.

40. Fan Z, Silva P, Gronau I, Wang S, Armero AS, Schweizer RM, et al. Worldwide patterns of genomic variation and admixture in gray wolves. Genome Res. 2016;26:163–73.

41. Chen H, Patterson N, Reich D. Population differentiation as a test for selective sweeps. Genome Res. 2010;20:393–402.

42. McLaren W, Gil L, Hunt SE, Riat HS, Ritchie GRS, Thormann A, et al. The Ensembl Variant Effect Predictor. Genome Biol. 2016;17:122.

43. Alexa A, Rahnenfuhrer J. topGO: Enrichment Analysis for Gene Ontology. 2016.

44. Grossmann S, Bauer S, Robinson PN, Vingron M. Improved detection of overrepresentation of Gene-Ontology annotations with parent child analysis. Bioinformatics. 2007;23:3024–31.

45. Maden M. Retinoic acid in the development, regeneration and maintenance of the nervous system. Nat. Rev. Neurosci. 2007;8:755–65.

46. Shirai H, Oishi K, Ishida N. Bidirectional CLOCK/BMAL1-dependent circadian gene regulation by retinoic acid in vitro. Biochem. Biophys. Res. Commun. 2006;351:387–91.

47. Slager RE, Newton TL, Vlangos CN, Finucane B, Elsea SH. Mutations in RAI1 associated with Smith–Magenis syndrome. Nat. Genet. 2003;33:466–466.

48. Potocki L, Bi W, Treadwell-Deering D, Carvalho CMB, Eifert A, Friedman EM, et al. Characterization of Potocki-Lupski syndrome (dup(17)(p11.2p11.2)) and delineation of a dosage-sensitive critical interval that can convey an autism phenotype. Am. J. Hum. Genet. 2007;80:633–633.

49. Olivares AM, Han Y, Soto D, Flattery K, Marini J, Mollema N, et al. The nuclear hormone receptor gene Nr2c1 (Tr2) is a critical regulator of early retina cell patterning. Dev. Biol. 2017;429:343–343.

50. Dedhar S, Rennie PS, Shago M, Hagesteijn CY, Yang H, Filmus J, et al. Inhibition of nuclear hormone receptor activity by calreticulin. Nature. 1994;367:480–480.

51. Takahashi H, Kanno T, Nakayamada S, Hirahara K, Sciumè G, Muljo SA, et al. TGF-β and retinoic acid induce the microRNA miR-10a, which targets Bcl-6 and constrains the plasticity of helper T cells. Nat. Immunol. 2012;13:587–587.

52. Yang J, Seo J, Nair R, Han S, Jang S, Kim K, et al. DGKé regulates presynaptic release during mGluR-dependent LTD. EMBO J. 2011;30:165–165.

53. Puranam RS, Eubanks JH, Heinemann SF, McNamara JO. Chromosomal localization of gene for human glutamate receptor subunit-7. Somat. Cell Mol. Genet. 1993;19:581–581.

54. Augustin I, Rosenmund C, Südhof TC, Brose N. Munc13-1 is essential for fusion competence of glutamatergic synaptic vesicles. Nature. 1999;400:457–457.

55. Caddick SJ, Wang C, Fletcher CF, Jenkins NA, Copeland NG, Hosford DA. Excitatory But Not Inhibitory Synaptic Transmission Is Reduced in Lethargic (Cacnb4 lh) and Tottering (Cacna1a tg) Mouse Thalami. J. Neurophysiol. 1999;81:2066–2066.

56. Harris JA, Westbrook RF. Evidence that GABA transmission mediates context-specific extinction of learned fear. Psychopharmacology. 1998;140:105–105.

57. Stork O, Ji FY, Obata K. Reduction of extracellular GABA in the mouse amygdala during and following confrontation with a conditioned fear stimulus. Neurosci. Lett. 2002;327:138–138.

58. Gassmann M, Bettler B. Regulation of neuronal GABAB receptor functions by subunit composition. Nat. Rev. Neurosci. 2012;13:380–380.

59. Oury F, Khrimian L, Denny CA, Gardin A, Chamouni A, Goeden N, et al. Maternal and offspring pools of osteocalcin influence brain development and functions. Cell. 2013;155:228–41.

60. Cheng L, Arata A, Mizuguchi R, Qian Y, Karunaratne A, Gray PA, et al. Tlx3 and Tlx1 are post-mitotic selector genes determining glutamatergic over GABAergic cell fates. Nat. Neurosci. 2004;7:510–510.

61. van Koningsbruggen S, Straasheijm KR, Sterrenburg E, de Graaf N, Dauwerse HG, Frants RR, et al. FRG1P-mediated aggregation of proteins involved in pre-mRNA processing. Chromosoma. 2007;116:53–53.

62. Mathew R, Hartmuth K, Möhlmann S, Urlaub H, Ficner R, Lührmann R. Phosphorylation of human PRP28 by SRPK2 is required for integration of the U4/U6-U5 tri-snRNP into the spliceosome. Nat. Struct. Mol. Biol. 2008;15:435–435.

63. Xia H, Chen D, Wu Q, Wu G, Zhou Y, Zhang Y, et al. CELF1 preferentially binds to exon-intron boundary and regulates alternative splicing in HeLa cells. Biochim. Biophys. Acta. 2017;1860:911–911.

64. Kim Y-D, Lee J-Y, Oh K-M, Araki M, Araki K, Yamamura K-I, et al. NSrp70 is a novel nuclear speckle-related protein that modulates alternative pre-mRNA splicing in vivo. Nucleic Acids Res. 2011;39:4300–4300.

65. Kim C-H, Kim Y-D, Choi E-K, Kim H-R, Na B-R, Im S-H, et al. Nuclear Speckle-related Protein 70 Binds to Serine/Arginine-rich Splicing Factors 1 and 2 via an Arginine/Serine-like Region and Counteracts Their Alternative Splicing Activity. J. Biol. Chem. 2016;291:6169–6169.

66. Straub T, Grue P, Uhse A, Lisby M, Knudsen BR, Tange TO, et al. The RNA-splicing factor PSF/p54 controls DNA-topoisomerase I activity by a direct interaction. J. Biol. Chem. 1998;273:26261–26261.

67. Lamers F, van der Ploeg I, Schild L, Ebus ME, Koster J, Hansen BR, et al. Knockdown of survivin (BIRC5) causes apoptosis in neuroblastoma via mitotic catastrophe. Endocr. Relat. Cancer. 2011;18:657–657.

68. Silke J, Vaux DL. Two kinds of BIR-containing protein - inhibitors of apoptosis, or required for mitosis. J. Cell Sci. 2001;114:1821–1821.

69. Redon R, Ishikawa S, Fitch KR, Feuk L, Perry GH, Andrews TD, et al. Global variation in copy number in the human genome. Nature. 2006;444:444–444.

70. Zhou Z, Jiang Y, Wang Z, Gou Z, Lyu J, Li W, et al. Resequencing 302 wild and cultivated accessions identifies genes related to domestication and improvement in soybean. Nat. Biotechnol. 2015;33:408–408.

71. Frantz LAF, Mullin VE, Pionnier-Capitan M, Lebrasseur O, Ollivier M, Perri A, et al. Genomic and archaeological evidence suggest a dual origin of domestic dogs. Science. 2016;352:1228–31.

72. Schlamp F, van der Made J, Stambler R, Chesebrough L, Boyko AR, Messer PW. Evaluating the performance of selection scans to detect selective sweeps in domestic dogs. Mol. Ecol. 2016;25:342–342.

73. Jensen JD. On the unfounded enthusiasm for soft selective sweeps [Internet]. 2014. Available from: http://dx.doi.org/10.1101/009563

74. Gotz S, Garcia-Gomez JM, Terol J, Williams TD, Nagaraj SH, Nueda MJ, et al. High-throughput functional annotation and data mining with the Blast2GO suite. Nucleic Acids Res. 2008;36:3420–3420.

75. Qanbari S, Pausch H, Jansen S, Somel M, Strom TM, Fries R, et al. Classic Selective Sweeps Revealed by Massive Sequencing in Cattle. PLoS Genet. 2014;10:e1004148.

76. Kijas JW. Haplotype-based analysis of selective sweeps in sheep. Genome. 2014;57:433–7.

77. Peyrégne S, Boyle MJ, Dannemann M, Prüfer K. Detecting ancient positive selection in humans using extended lineage sorting. Genome Res. 2017;27:1563–1563.

78. Prüfer K, Racimo F, Patterson N, Jay F, Sankararaman S, Sawyer S, et al. The complete genome sequence of a Neanderthal from the Altai Mountains. Nature. 2014;505:43–43.

79. Racimo F. Testing for Ancient Selection Using Cross-population Allele Frequency Differentiation. Genetics. 2016;202:733–733.

80. Lin R, Du X, Peng S, Yang L, Ma Y, Gong Y, et al. Discovering All Transcriptome Single-Nucleotide Polymorphisms and Scanning for Selection Signatures in Ducks (Anas platyrhynchos). Evol. Bioinform. Online. 2015;11:67–67.

81. Schubert M, Jónsson H, Chang D, Der Sarkissian C, Ermini L, Ginolhac A, et al. Prehistoric genomes reveal the genetic foundation and cost of horse domestication. Proc. Natl. Acad. Sci. U. S. A. 2014;111:E5661–5661.

82. Sánchez-Villagra MR, Geiger M, Schneider RA. The taming of the neural crest: a developmental perspective on the origins of morphological covariation in domesticated mammals. R Soc Open Sci. 2016;3:160107.

83. Bronner ME, LeDouarin NM. Development and evolution of the neural crest: An overview. Dev. Biol. 2012;366:2–2.

84. van Es JH, Haegebarth A, Kujala P, Itzkovitz S, Koo B-K, Boj SF, et al. A critical role for the Wnt effector Tcf4 in adult intestinal homeostatic self-renewal. Mol. Cell. Biol. 2012;32:1918–1918.

85. Tang L-Y, Deng N, Wang L-S, Dai J, Wang Z-L, Jiang X-S, et al. Quantitative phosphoproteome profiling of Wnt3a-mediated signaling network: indicating the involvement of ribonucleoside-diphosphate reductase M2 subunit phosphorylation at residue serine 20 in canonical Wnt signal transduction. Mol. Cell. Proteomics. 2007;6:1952–1952.

86. Bergmann C, Fliegauf M, Brüchle NO, Frank V, Olbrich H, Kirschner J, et al. Loss of nephrocystin-3 function can cause embryonic lethality, Meckel-Gruber-like syndrome, situs inversus, and renal-hepatic-pancreatic dysplasia. Am. J. Hum. Genet. 2008;82:959–959.

87. Carmon KS, Gong X, Lin Q, Thomas A, Liu Q. R-spondins function as ligands of the orphan receptors LGR4 and LGR5 to regulate Wnt/beta-catenin signaling. Proc. Natl. Acad. Sci. U. S. A. 2011;108:11452–11452.

88. Bronner-Fraser M. Neural crest cell formation and migration in the developing embryo. FASEB J. 1994;8:699–699.

89. Santagati F, Rijli FM. Cranial neural crest and the building of the vertebrate head. Nat. Rev. Neurosci. 2003;4:806–806.

90. Lagadec R, Laguerre L, Menuet A, Amara A, Rocancourt C, Péricard P, et al. The ancestral role of nodal signalling in breaking L/R symmetry in the vertebrate forebrain. Nat. Commun. 2015;6:6686.

91. Berryman MA, Goldenring JR. CLIC4 is enriched at cell-cell junctions and colocalizes with AKAP350 at the centrosome and midbody of cultured mammalian cells. Cell Motil. Cytoskeleton. 2003;56:159–159.

92. de la Torre-Ubieta L, Gaudilliére B, Yang Y, Ikeuchi Y, Yamada T, DiBacco S, et al. A FOXO-Pak1 transcriptional pathway controls neuronal polarity. Genes Dev. 2010;24:799–799.

93. Kunisaki Y, Nishikimi A, Tanaka Y, Takii R, Noda M, Inayoshi A, et al. DOCK2 is a Rac activator that regulates motility and polarity during neutrophil chemotaxis. J. Cell Biol. 2006;174:647–647.

94. Dupraz S, Grassi D, Bernis ME, Sosa L, Bisbal M, Gastaldi L, et al. The TC10-Exo70 complex is essential for membrane expansion and axonal specification in developing neurons. J. Neurosci. 2009;29:13292–13292.

95. Le Douarin NM, Dupin E. The neural crest in vertebrate evolution. Curr. Opin. Genet. Dev. 2012;22:381–381.

96. Minoux M, Rijli FM. Molecular mechanisms of cranial neural crest cell migration and patterning in craniofacial development. Development. 2010;137:2605–2605.

97. Morey DF. Size, shape and development in the evolution of the domestic dog. J. Archaeol. Sci. 1992;19:181–181.

98. Etchevers HC, Couly G, Vincent C, Le Douarin NM. Anterior cephalic neural crest is required for forebrain viability. Development. 1999;126:3533–3533.

99. Xavier GM, Sharpe PT, Cobourne MT. Scube1 is expressed during facial development in the mouse. J. Exp. Zool. B Mol. Dev. Evol. 2009;312B:518–518.

100. FitzPatrick DR, Carr IM, McLaren L, Leek JP, Wightman P, Williamson K, et al. Identification of SATB2 as the cleft palate gene on 2q32-q33. Hum. Mol. Genet. 2003;12:2491–501.

101. Dobreva G, Chahrour M, Dautzenberg M, Chirivella L, Kanzler B, Fariñas I, et al. SATB2 is a multifunctional determinant of craniofacial patterning and osteoblast differentiation. Cell. 2006;125:971–971.

102. Szeto DP, Rodriguez-Esteban C, Ryan AK, O’Connell SM, Liu F, Kioussi C, et al. Role of the Bicoid-related homeodomain factor Pitx1 in specifying hindlimb morphogenesis and pituitary development. Genes Dev. 1999;13:484–484.

103. St Amand TR, Zhang Y, Semina EV, Zhao X, Hu Y, Nguyen L, et al. Antagonistic signals between BMP4 and FGF8 define the expression of Pitx1 and Pitx2 in mouse tooth-forming anlage. Dev. Biol. 2000;217:323–323.

104. Rapini RP, Warner NB. Relapsing polychondritis. Clin. Dermatol. 2006;24:482–482.

105. Boyko AR, Quignon P, Li L, Schoenebeck JJ, Degenhardt JD, Lohmueller KE, et al. A simple genetic architecture underlies morphological variation in dogs. PLoS Biol. 2010;8:e1000451.

106. Nagai N, Hosokawa M, Itohara S, Adachi E, Matsushita T, Hosokawa N, et al. Embryonic lethality of molecular chaperone hsp47 knockout mice is associated with defects in collagen biosynthesis. J. Cell Biol. 2000;150:1499–1499.

107. Wilson R, Norris EL, Brachvogel B, Angelucci C, Zivkovic S, Gordon L, et al. Changes in the chondrocyte and extracellular matrix proteome during post-natal mouse cartilage development. Mol. Cell. Proteomics. 2012;11:M111.014159.

108. Argente J, Flores R, Gutiérrez-Arumí A, Verma B, Martos-Moreno GÁ, Cuscó I, et al. Defective minor spliceosome mRNA processing results in isolated familial growth hormone deficiency. EMBO Mol. Med. 2014;6:299–299.

109. An M, Henion PD. The zebrafish sf3b1b460 mutant reveals differential requirements for the sf3b1 pre-mRNA processing gene during neural crest development. Int. J. Dev. Biol. 2012;56:223–223.

110. Edery P, Marcaillou C, Sahbatou M, Labalme A, Chastang J, Touraine R, et al. Association of TALS developmental disorder with defect in minor splicing component U4atac snRNA. Science. 2011;332:240–240.

111. Agnvall B, Jöngren M, Strandberg E, Jensen P. Heritability and genetic correlations of fear-related behaviour in Red Junglefowl possible implications for early domestication. PLoS One. 2012;7:e35162.

112. Lindberg J, Björnerfeldt S, Saetre P, Svartberg K, Seehuus B, Bakken M, et al. Selection for tameness has changed brain gene expression in silver foxes. Curr. Biol. 2005;15:R915–915.

113. Trut LN, Plyusnina IZ, Oskina IN. An Experiment on Fox Domestication and Debatable Issues of Evolution of the Dog. Russ. J. Genet. 2004;40:644–644.

114. Hare B, Wobber V, Wrangham R. The self-domestication hypothesis: evolution of bonobo psychology is due to selection against aggression. Anim. Behav. 2012;83:573–573.

115. Morey DF, Jeger R. Paleolithic dogs: Why sustained domestication then? Journal of Archaeological Science: Reports. 2015;3:420–420.

116. Li Y, Wang G-D, Wang M-S, Irwin DM, Wu D-D, Zhang Y-P. Domestication of the dog from the wolf was promoted by enhanced excitatory synaptic plasticity: a hypothesis. Genome Biol. Evol. 2014;6:3115–3115.

117. Truong HT, Solaymani-Kohal S, Baker KR, Girirajan S, Williams SR, Vlangos CN, et al. Diagnosing Smith-Magenis syndrome and duplication 17p11.2 syndrome by RAI1 gene copy number variation using quantitative real-time PCR. Genet. Test. 2008;12:67–67.

118. Elsea SH, Girirajan S. Smith–Magenis syndrome. Eur. J. Hum. Genet. 2008;16:412–412.

119. vonHoldt BM, Shuldiner E, Koch IJ, Kartzinel RY, Hogan A, Brubaker L, et al. Structural variants in genes associated with human Williams-Beuren syndrome underlie stereotypical hypersociability in domestic dogs. Sci Adv. 2017;3:e1700398.

120. Jones W, Bellugi U, Lai Z, Chiles M, Reilly J, Lincoln A, et al. II. Hypersociability in Williams Syndrome. J. Cogn. Neurosci. 2000;12 Suppl 1:30–30.

121. Adams MS, Gammill LS, Bronner-Fraser M. Discovery of transcription factors and other candidate regulators of neural crest development. Dev. Dyn. 2008;237:1021–1021.

122. Tahir R, Kennedy A, Elsea SH, Dickinson AJ. Retinoic acid induced-1 (Rai1) regulates craniofacial and brain development in Xenopus. Mech. Dev. 2014;133:91–91.

123. Barnett C, Yazgan O, Kuo H-C, Malakar S, Thomas T, Fitzgerald A, et al. Williams Syndrome Transcription Factor is critical for neural crest cell function in Xenopus laevis. Mech. Dev. 2012;129:324–324.

124. Goldman SE, Malow BA, Newman KD, Roof E, Dykens EM. Sleep patterns and daytime sleepiness in adolescents and young adults with Williams syndrome. J. Intellect. Disabil. Res. 2009;53:182–182.

125. Sniecinska-Cooper AM, Iles RK, Butler SA, Jones H, Bayford R, Dimitriou D. Abnormal secretion of melatonin and cortisol in relation to sleep disturbances in children with Williams syndrome. Sleep Med. 2015;16:94–94.

126. Williams SR, Zies D, Mullegama SV, Grotewiel MS, Elsea SH. Smith-Magenis syndrome results in disruption of CLOCK gene transcription and reveals an integral role for RAI1 in the maintenance of circadian rhythmicity. Am. J. Hum. Genet. 2012;90:941–941.

127. De Leersnyder H, De Blois MC, Claustrat B, Romana S, Albrecht U, Von Kleist-Retzow JC, et al. Inversion of the circadian rhythm of melatonin in the Smith-Magenis syndrome. J. Pediatr. 2001;139:111–111.

128. Kolesnikova LA. [Circadian rhythm of biosynthetic activity of the epiphysis in relatively wild and domesticated silver foxes]. Genetika. 1997;33:1144–1144.

129. Popova NK, Kulikov AV, Avgustinovich DF, Voĭtenko NN, Trut LN. [Effect of domestication of the silver fox on the main enzymes of serotonin metabolism and serotonin receptors]. Genetika. 1997;33:370–370.

130. Popova NK, Voitenko NN, Kulikov AV, Avgustinovich DF. Evidence for the involvement of central serotonin in mechanism of domestication of silver foxes. Pharmacol. Biochem. Behav. 1991;40:751–751.

131. McKenna A, Hanna M, Banks E, Sivachenko A, Cibulskis K, Kernytsky A, et al. The Genome Analysis Toolkit: a MapReduce framework for analyzing next-generation DNA sequencing data. Genome Res. 2010;20:1297–1297.

132. Alexander DH, Novembre J, Lange K. Fast model-based estimation of ancestry in unrelated individuals. Genome Res. 2009;19:1655–1655.

133. Purcell S, Neale B, Todd-Brown K, Thomas L, Ferreira MAR, Bender D, et al. PLINK: a tool set for whole-genome association and population-based linkage analyses. Am. J. Hum. Genet. 2007;81:559–559.

134. Patterson N, Price AL, Reich D. Population structure and eigenanalysis. PLoS Genet. 2006;2:e190.

135. Purcell S, Neale B, Todd-Brown K, Thomas L, Ferreira MAR, Bender D, et al. PLINK: a tool set for whole-genome association and population-based linkage analyses. Am. J. Hum. Genet. 2007;81:559–559.

136. Danecek P, Auton A, Abecasis G, Albers CA, Banks E, DePristo MA, et al. The variant call format and VCFtools. Bioinformatics. 2011;27:2156–2156.

137. Kelleher J, Etheridge AM, McVean G. Efficient Coalescent Simulation and Genealogical Analysis for Large Sample Sizes. PLoS Comput. Biol. 2016;12:e1004842.

138. Campbell CL, Bhérer C, Morrow BE, Boyko AR, Auton A. A Pedigree-Based Map of Recombination in the Domestic Dog Genome. G3 [Internet]. 2016; Available from: http://dx.doi.org/10.1534/g3.116.034678

139. Bhatia G, Patterson N, Sankararaman S, Price AL. Estimating and interpreting FST: The impact of rare variants. Genome Res. 2013;23:1514–1514.

140. Price AL, Patterson NJ, Plenge RM, Weinblatt ME, Shadick NA, Reich D. Principal components analysis corrects for stratification in genome-wide association studies. Nat. Genet. nature.com; 2006;38:904–904.

141. Pavlidis P, Noble WS. Matrix2png: a utility for visualizing matrix data. Bioinformatics. Oxford Univ Press; 2003;19:295–295.

142. Gotz S, Garcia-Gomez JM, Terol J, Williams TD, Nagaraj SH, Nueda MJ, et al. High-throughput functional annotation and data mining with the Blast2GO suite. Nucleic Acids Res. 2008;36:3420–3420.

143. Ramirez O, Olalde I, Berglund J, Lorente-Galdos B, Hernandez-Rodriguez J, Quilez J, et al. Analysis of structural diversity in wolf-like canids reveals post-domestication variants. BMC Genomics. 2014;15:465.

